# Long-term development of a motor memory

**DOI:** 10.1101/2025.02.22.639647

**Authors:** David W. Franklin, Pascal Nietschmann, Thalia Papadopoulou, Sae Franklin

## Abstract

Human behavior is developed through continuous adaptation to our environment over a range of timescales. Extensive studies have investigated the mechanisms and computations underlying this process of sensorimotor adaptation using several hundred trials. However, most of our motor skills have had countless hours of practice. Here we study a simple motor adaptation task using thousands of training trials over multiple weeks to study the long-term development of a motor memory, and examine changes in adaptation, retention, inter-limb transfer, decay, spontaneous recovery and generalization. Unlike previous studies, participants showed complete compensation to the novel dynamics, along with long-term increases in retention and spontaneous recovery. Moreover, we find narrowing in the angular generalization, suggesting continual tuning of the motor memory to the task. This demonstrates the extensive changes occurring with longer training of motor tasks, highlighting their importance in studies of sensorimotor control, rehabilitation and training.

## Introduction

Human behavior has kept evolving throughout our history. As new tools are developed, we continually adapt to accommodate them. We are now able to perform a wide variety of skillful actions in our daily life, continuously interacting with our environment, all driven by our complex sensorimotor control system ^1^. Such skill learning requires the development and retainment of motor memories of these tasks, where these motor memories incorporate time varying patterns of control processes within a motion, and the sequencing of these motions to produce a complex task. This large repertoire of motor skills is only possible by the process of adaptation and generalization, in which motor memories are formed, adjusted, or combined for each new action ^2–4^. Understanding the mechanisms underlying the development of motor memories is not only critical to understand sensorimotor control ^5–7^, but it also is necessary for improving learning and rehabilitation ^8,9^. To understand the mechanisms underlying the development of a motor memory, studies have investigated adaptation to novel dynamics ^10–16^ or novel sensorimotor transformations ^17–20^ using controlled designs in which the trial-by-trial motor adaptation can be examined. Adaptation to novel dynamics normally occurs rapidly using sensed error signals (proprioceptive and visual error information) to update the current motor memory. Repeated adaptation leads to further updates of our internal models and a more permanent motor memory ^21,22^. Once the internal model is acquired, it then becomes available for recall with the appropriate contextual cues ^3,23–25^.

To investigate in detail the mechanisms which drive motor adaptation, studies employ controlled novel manipulations in the laboratory, in which specific patterns of trials involving force fields, novel sensorimotor transformations, channels, error clamps, and probes are presented to quantitatively assess measures of learning in the temporal domain ^12–15,26–30^. These studies have unveiled the computational mechanisms that underlie motor memory formation, selection, generalization and representation ^2,4,7,23,31,32^. However, most motor adaptation studies examine only the changes within one or two hours of practice.

In contrast, most of our developed skills are performed many times each day, leading to thousands and thousands of repetitions over years. Development of skills over months or years has been examined in tasks such as cigar making, mirror tracing, and visual tracking tasks ^33^. Repeated training improves their skills over months and years ^34^ leading to long term retention and permanent skill acquisition ^35,36^. How does repeated training affect the formation, selection and transfer of motor memories? Adaptation to visuomotor rotations strengthening effects over several sessions ^37^. Only a few studies have examined longer-term acquisition of novel dynamics in a laboratory setting, often using non-human primates ^38^ or complex manipulations ^39^. We have many remaining open questions regarding how training effects motor memories, such as its effect on retention, cross-limb transfer, or representation. To address this, here we investigate the effect of long-term practice of a simple motor adaptation task over 8 consecutive weeks with healthy human participants, testing how extensive practice effects the retention, decay, transfer, and generalization of the motor memories. Finally, we performed a full recall session, 15 weeks after the last training session, to investigate how the memory is influenced by long periods with no practice.

## Results

Ten participants performed reaching movements over 8 weeks of training followed by a final session 15 weeks later (week 23). After an initial familiarization phase performed in the null field, the participants were introduced to the curl force field. Although each session contained different trials to assess the learning, retention, transfer, and generalization of the learned motor memory, the participants generally continued to perform the same forward reaching movement in the curl force field for the rest of the sessions, with over 7000 trials.

The kinematic error, which was low in the initial null force field trials, increased upon initial exposure to the CCW curl field (Fig 1A). This reduced over the course of the first session and continued to reduce over the subsequent training sessions. In weeks 1, 7 and 23, the opposite curl force field (CW field) was applied for 30 trials, which produced large kinematic errors in the opposite direction. Force compensation over the full experiment showed similar performance increases over the weeks of training (Fig 1B). The force compensation reached about 80% by the end of the first training session, continued to increase over the following weeks plateauing over 90% by the end of the experiments. The effects of introducing both the opposing curl force field and repeated channel trials can also been seen in the force compensation over the different weeks, with decreases in the force compensation. Both reaction time (Fig 1C) and peak speed (Fig 1D) were consistent across the training weeks. The mean peak speed quickly aligned to the middle of the “great” feedback zone and stayed near that level throughout the experiment, with only noticeable reductions during the introduction of the opposite curl force field (red traces).

**Figure 1.**
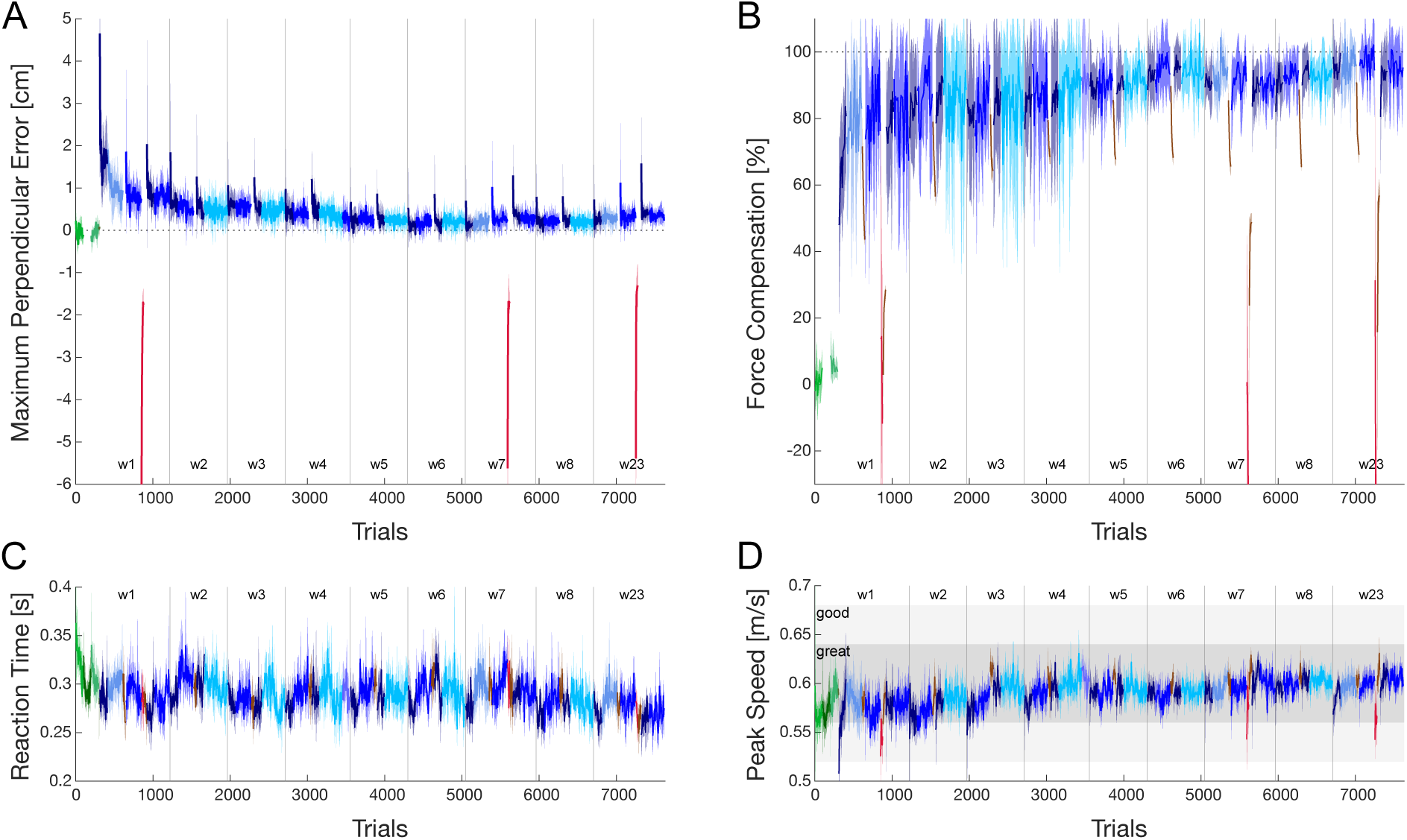
Adaptation over 9 sessions of training. **A**. Mean kinematic error across participants as measured by the maximum perpendicular error in each right-hand reaching movement. Here positive errors are shown for the effects of the main force field used (CCW), that is for errors in the negative x-axis. The color indicates the condition of the experiment (see Methods). Overall, green indicates movements in the null force field, blue and cyan indicate movements in the curl force field (CCW), red indicates movements in the opposite curl force field (CW), and brown indicates phases of channel trials. The dotted lines indicate the period between each session. The line indicates the mean and the shaded region indicates the standard error of the mean across all participants. **B**. Mean force compensation across participants expressed as a percentage of perfect compensation to the force field measured during right-hand channel trials in the forward direction throughout the experiment. Colors indicate the condition of the experiment in which the values were measured. **C**. Mean reaction time throughout the experiment. **D**. Mean peak speed throughout the experiment. The light grey region indicates peaks speeds in which participants received “good” feedback on their speed, whereas the dark grey region indicates speeds for which participants received “great” feedback.

Although the kinematic error showed a strong reduction in the size of the maximum error over the experiment, it only reflects one time point in each trajectory. To more fully examine the path over which participants were moving we plotted the full trajectories at specific points in the sessions. As expected, the mean trajectory in the null field was straight to the target (Fig 2A), but the initial trajectories in the curl force field exhibited strong disturbances to the left. Consecutive trajectories show a reduction in the size of the error, but this is quickly reduced, such that the tenth trial is not much straighter than the fifth trial, and the final trajectories at the end of week 1 are only slightly straighter. However, it can also be seen that the final trajectories on week 8 look much closer to those in the null field. To examine this further we plotted the final trajectories at the end of each training day with a zoomed in x-axis (Fig 2B). Although the mean trajectory at the end of the first week’s session (w1, light blue) was already reduced dramatically from the initial perturbed trajectories, it still has an error, peaking around 8 mm laterally. However, it is clearly seen that further training weeks result in straighter and straighter movements. Interestingly, the s-shape that is seen in the first few weeks never disappears from the mean trajectories. Even the final sessions (w8 and w23) show subtle s-shape trajectories even though the maximum lateral errors from the straight line are only around 1-2 mm. These values are similar to those in the null field (green line) but follow a very different trajectory. These S-shaped hand paths have previously been shown when participants adapted to a velocity dependent curl force field. This shape is created when hand paths tend to overcompensate the field early in the movement and slightly undercompensate when approaching the target, which has been suggested to indicate that participants optimize control by reducing motor costs ^40^.

**Figure 2.**
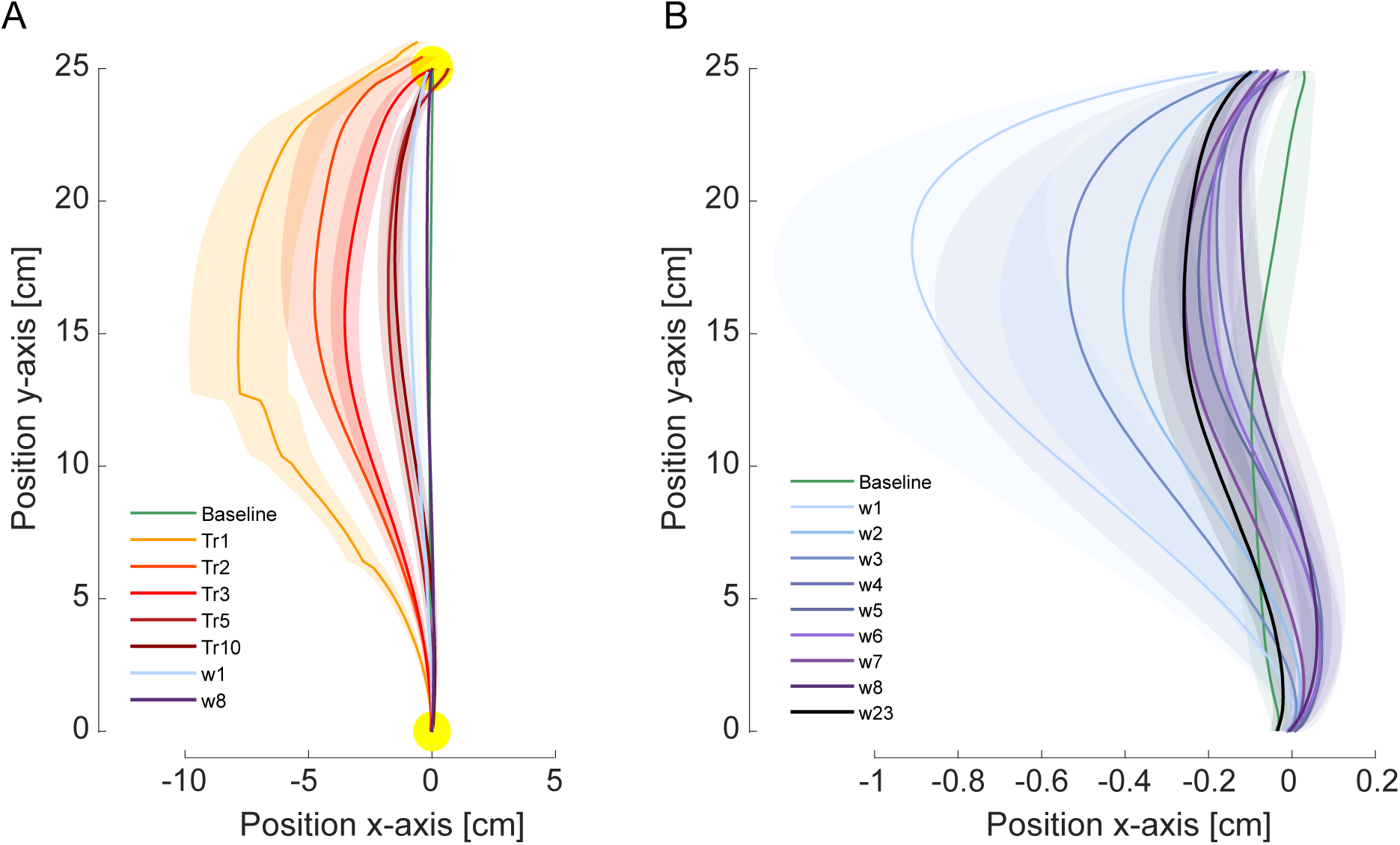
Movement trajectories over the training sessions. **A**. Trajectories in the initial session of training (week 1). Each line indicates the mean trajectory across the 10 participants, with the shaded region indicating the standard error of the mean. Baseline indicates the final trial in the null field prior to the introduction of the curl force field, whereas Tr1-Tr10 indicate the first, second, third, fifth and tenth trials in the curl field. The final curl field trajectories at the end of week 1 and week 8 are shown for comparison. **B**. The final trajectories at the end of each week’s session are shown across the experiment. Each line indicates the mean across participants of the final trial in each week’s session. Baseline indicates the final trial in the null field in week 1.

The kinematic trajectories suggest that adaptation has proceeded further with additional training sessions over the weeks. However, as kinematic error can be affected both by feedforward adaptation and by increased mechanical impedance due to co-contraction, we examined the pattern of predictive force compensation on the channel trials at the end of learning in each week’s session. While the measure of force compensation provides a measure of the predictive compensation it is still possible to obtain a force compensation of near 100% when the predictive force does not match the temporal pattern required for compensation. Therefore, in order to examine the time varying pattern of force compensation, we aligned the individual forward force and velocity traces to the movement start, velocity peak, and movement end of each trial before averaging. This allows us to both average across multiple trials and multiple participants without affecting the force pattern. The force pattern on the channel trials over the sessions can be seen relative to the forces on the curl force field trials (Fig 3A). Although the predictive force profiles on the channel trials are close to that of the forces in the curl field after training in the first week, there are still differences that persist for the next few sessions. Specifically, we see a high variability across participants for the first three weeks. However, by the fourth and subsequent weeks these two force profiles match almost perfectly. To quantify the exact values, we calculate the predictive compensation, which shows that for weeks 4 to 8, the predictive compensation is over 95% across all ten participants (Fig 3B). A repeated measures ANOVA found no significant differences across the 9 sessions (F_1.563,14.063_=1.261; p=0.304; 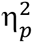=0.123). Overall, we see that with further training in the curl force field, the predictive compensation reaches over 95% and matches almost perfectly the forces experienced in the force field.

**Figure 3.**
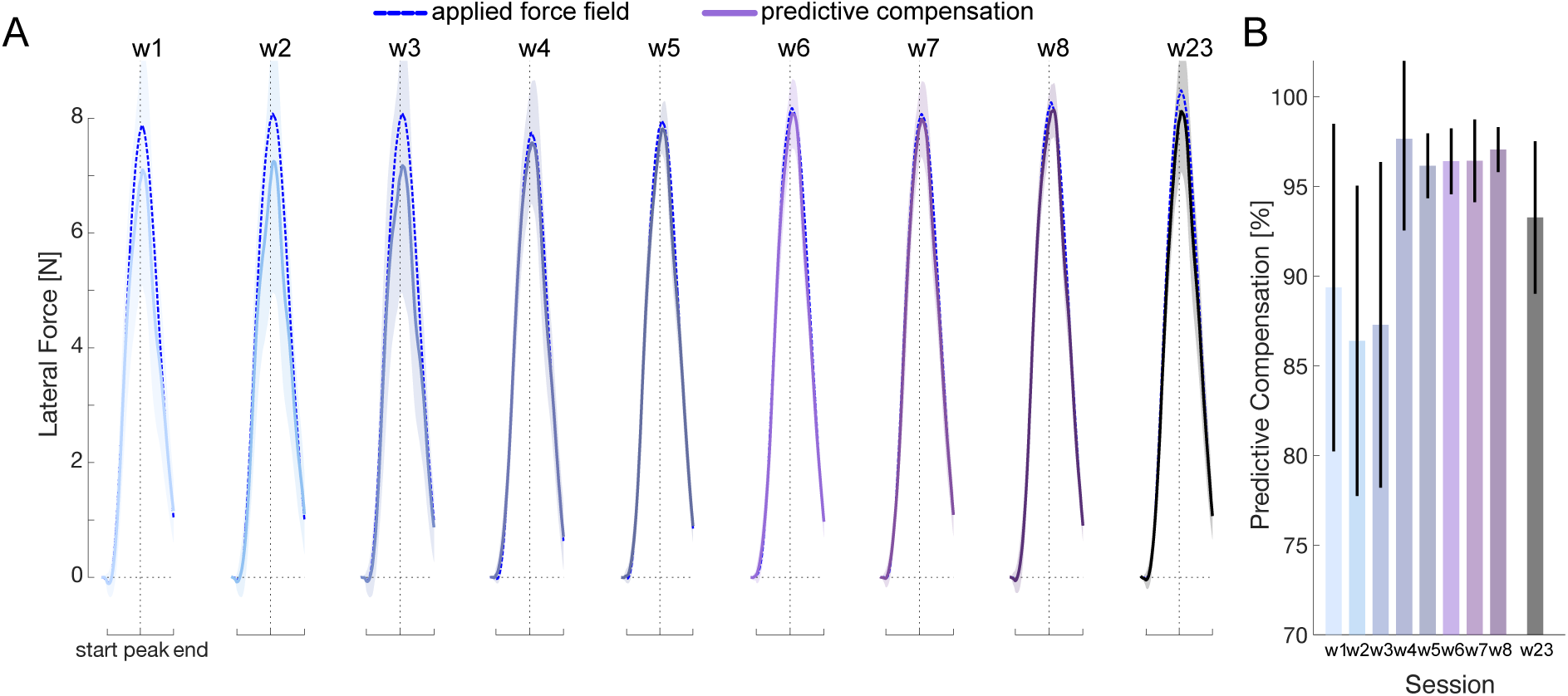
Predictive force compensation at the end of each training session. **A**. The predictive force exerted against the channel wall (solid line) and the force produced by the force field (dashed line) at the end of each week’s session. Each line represents the mean across 10 trials for each of the participants. The shaded region indicates the standard error of the mean. Prior to averaging the force traces were aligned to the start, velocity peak and end of each trial. The dotted black line shows the time of the peak velocity. **B**. The percent of predictive compensation seen on the channel trials. The error bar indicates the standard error of the mean.

### Motor Memory Generalization

To examine the generalization of the learned motor memory to similar reaching directions, we introduced random reaching movements to six other directions in channel trials throughout the training sessions (see Methods). For each week’s session the force compensation on the channel trials was calculated and plotted as a function of the movement angle (Fig 4A). The force compensation prior to the introduction of the force field (baseline, green trace) shows little to no forces across the different reaching directions as expected. By the end of the first session (w1, light blue trace), the peak of the force compensation was above 80% at the training direction and showed strong generalization to neighboring directions. However, in later training sessions both the peak of the force compensation appeared to increase, and the width of the generalization appeared to decrease (darker color traces). To quantify the degree of changes in the generalization we fit the data with Gaussian functions using a leave-two out bootstrapping method. It can be seen that over the training sessions the peak of the Gaussian increased significantly (based on 95% confidence intervals), from around 80% in the first few sessions to over 95% in the final w23 session (Fig 4B) and the width of the Gaussian decreased significantly, from around 30° in the first few sessions to 20° in the final w23 session (Fig 4C). Overall, this shows a narrowing of the generalization with adaptation as the motor memory is further trained.

**Figure 4.**
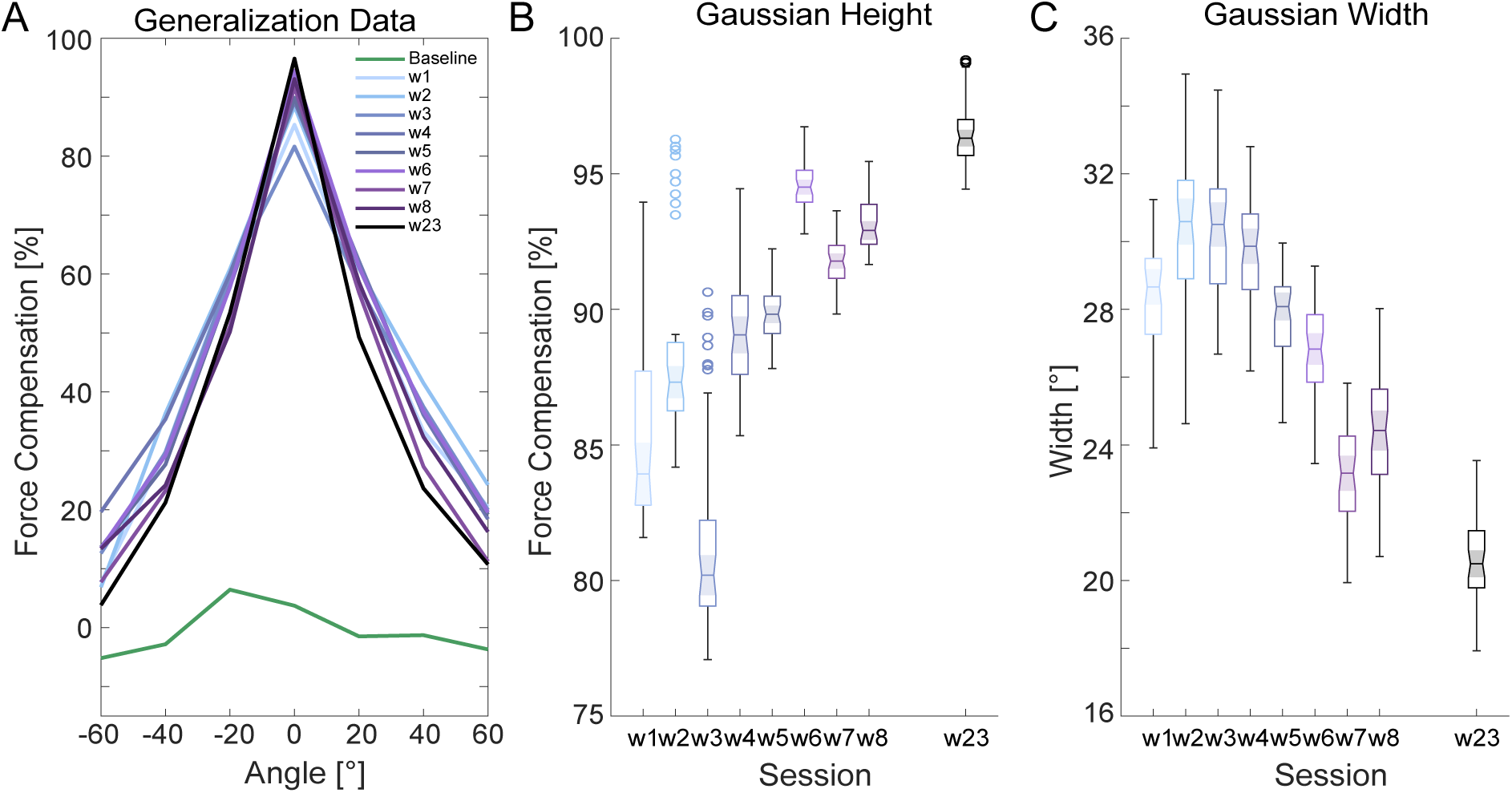
Generalization of the learned motor memory to different reaching angles. **A**. Force compensation expressed on reaching movements to targets at different angles (−60° to +60°, where 0° indicates the training direction). All measurements are made on channel trials. Note that participants were never exposed to the curl force field on the −60°, −40°, −20°, 20°, 40° or 60° directions. **B**. The best-fit parameters for the height of the Gaussian function over training sessions. Model parameters were obtained with the leave-two-out cross validation method, which provided 45 estimates of each parameter. Parameters estimates were plotted with boxchart in Matlab where the line indicates the mean, the shaded notch indicates the 95% confidence intervals, the upper and lower edges of the box contain the upper and lower quartiles, the whiskers contain the nonoutlier maximum and minimum, and any outliers are indicated with small circles. If the shaded notch regions do not overlap, then the parameters have different medians at the 5% significance level. **C**. The best-fit model parameters of the width of the Gaussian function.

### Motor Memory Decay

It has long been known that the motor memory decays in the absence of error information ^41^. This has been taken as evidence that the retention factor of the adaptation mechanism is less than 1.0, meaning that we forget a little part of what we have learned on each subsequent trial ^32,41^. It has been suggested that our motor adaptation system has this forgetting factor in order to continually search for minimum metabolic cost approaches to solving the current task ^7,42^. Recent work has suggested the existence of multiple timescales of adaptation, such as fast, slow or even hyperslow adaptation processes ^27,32,43,44^. In such a framework, we might expect that the decay of the learned motor memory could vary over longer time exposures to the changed dynamics. Here we examined the time course of the decay using 40 channel trials in a row during every training session. As expected, the force compensation on these channel trials decays over the consecutive trials (Fig 5A). However, the amount of decay is lower for later sessions than for the initial weeks. To quantify the rate of decay we fit an exponential function to the channel trial data (Fig 5B). Using a leave-two-out cross validation method we determined the best-fit parameters of the exponential function. Initial compensation level (Fig 5C) was determined using the last 5 channel trials during the force field applications. As expected from our previous results, this level increased over the training sessions, reaching around 95% in the later sessions. However, despite the increased adaptation level in the later sessions, the amount of decay decreased from the initial session (where it was approximately 50%) to around 30% in the later sessions (Fig 5D). The time constant of the decay (Fig 5D) was fairly consistent across the sessions, with the exception of week 8 where there was high variability. Overall, we found a reduction in the amount of decay but no consistent effect on the time course of the decay over sessions.

**Figure 5.**
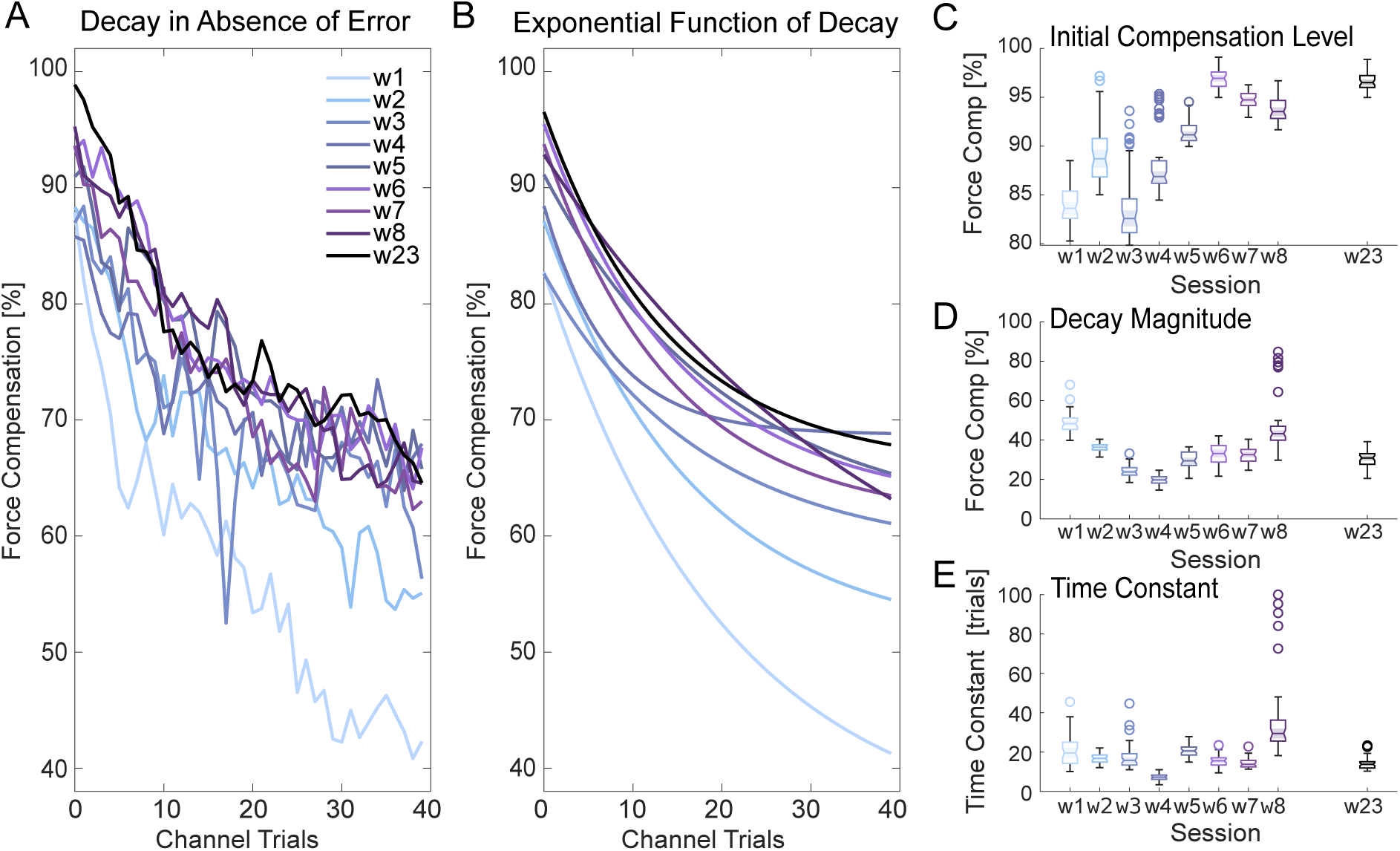
Trial-dependent decay of the motor memory in the absence of error information. **A**. The decay of force compensation in consecutive channel trials in which the lateral error is constrained to zero. Each trace illustrates the mean force compensation across the ten participants in the channel trials. Colors denote the specific sessions. **B**. Best-fit exponential functions to the mean force compensation across the different sessions. **C**. Initial force compensation level at the beginning of the repeated channel trials. Values are determined from last five channel trials in the prior force field phase. Parameters estimates were plotted with boxchart in Matlab where the line indicates the mean, the shaded notch indicates the 95% confidence intervals, the upper and lower edges of the box contain the upper and lower quartiles, the whiskers contain the nonoutlier maximum and minimum, and any outliers are indicated with small circles. If the shaded notch regions do not overlap, then the parameters have different medians at the 5% significance level. **D**. Magnitude of the decay as estimated from a best-fit exponential function of the force compensation. Model parameters were obtained with the leave-two-out cross validation method, which provided 45 estimates of each parameter. **E**. Time constant of the decay as estimated from a best-fit exponential function of the force compensation during the channel trials.

### Motor Memory Retention between Sessions

It has been shown that motor memories of the novel dynamics are retained over a long period of time in the absence of experience on the robotic systems ^21,22^. However, as these previous studies quantified the retention only by using the correlation coefficient of the velocity profiles between sessions, and did not actually measure the forces against channel walls, it is not clear to what degree the motor memories are specifically retained. For example, if participants simply co-contracted their muscles to increase the stiffness of their limbs ^11,45,46^, they could produce similar trajectories, resulting in a similar level of retention according to this metric. Here we investigate the retention of the learning between sessions using 5 channel trials at the beginning of each session prior to any force field trials. The retention of the learned force profile at the start of each session was compared with the final predictive force profiles on the previous session (Fig 6A). While the predictive force profile at the end of week 1 reached around 7N at its peak, the retained force profile on week two was less than 2N at its peak (Fig 6A, leftmost plot). However, there is still a rough shape of the profile, peaking at the time of peak velocity and close to zero force at the start and end of each movement. By week 3 the retained force profile is higher, reaching a peak of 4N with a temporal profile similar to the previous week. This retained force profile increases on the subsequent weeks, but with little changes beyond week 6. Therefore, even with a week between sessions, most of the learned predictive force profile is retained, with a clear temporal profile matching that of the previous session. Finally, in week 23, after 15 weeks with no experience of the force field, the retention was again measured with the channel trials (Fig 6A, rightmost plot), showing similar levels of retention as after only 1 week. To examine the amount of retention compared to the actual amount learned, we quantify the ratio of retention between the initial predictive force profile and the final force profile on the previous session (Fig 6B). We see that after week two, in which this retention is only around 20% (18.71%), the retention ranges between 58.15% (week 3) and 69.86% (week 8) with little differences between weeks 3 to 23. Using a repeated measures ANOVA we found a significant main effect of week (F_7,63_=7.089; p<0.001; 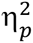=0.441). Bonferroni-corrected post-hoc comparisons showed that the retention on week 2 was significantly less than all other weeks (all p<0.021) except for week 7 (p=0.076). However, there were no differences across weeks 3 to 23 (all p=1.0). Overall, here we show that the learned predictive profiles are retained across multiple weeks, exhibiting almost slightly scaled down versions of the temporal profiles of the force, but maintaining even 60% retention after 15 weeks with no experience of the force fields.

**Figure 6.**
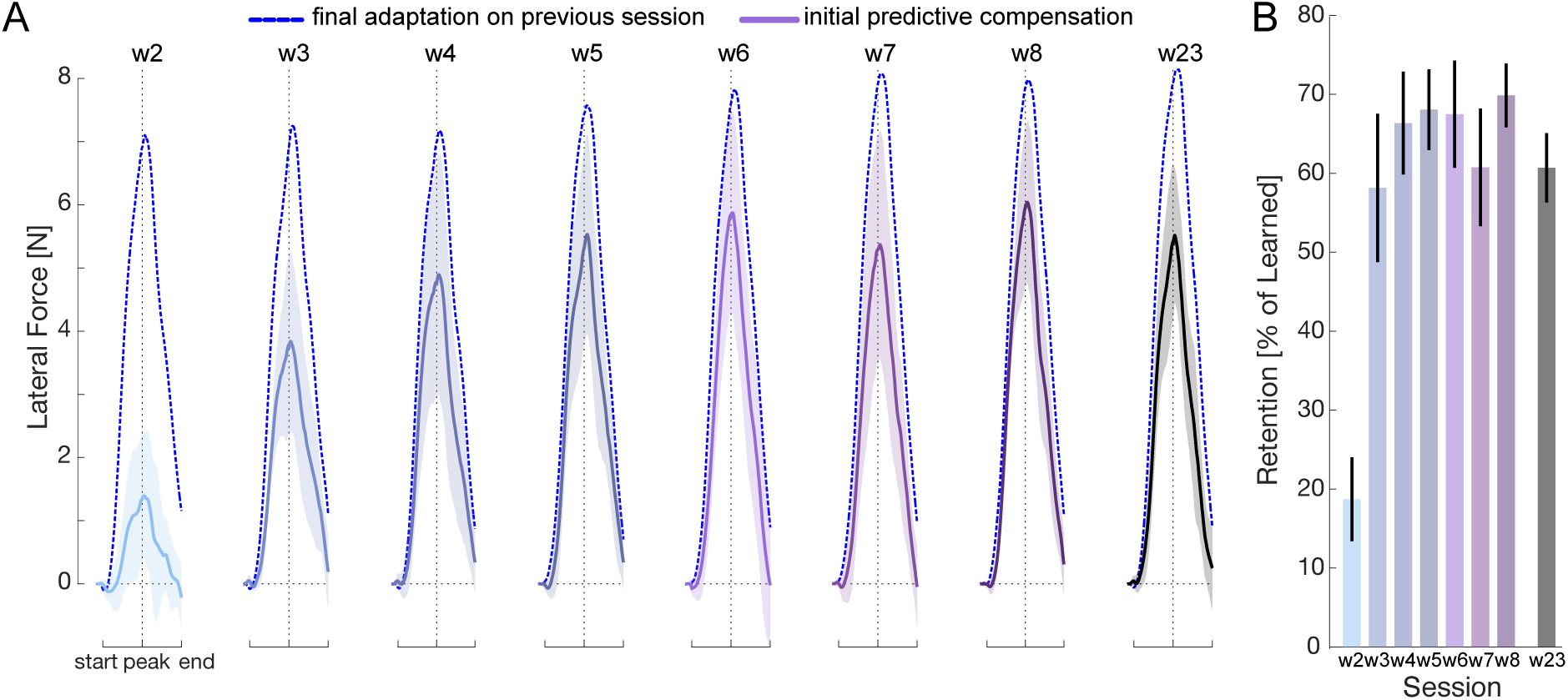
Retention of predictive force profile between sessions. **A**. The pattern of lateral force exerted against the channel wall during the first five trials at the beginning of each session (colored lines) compared to the forces measured during random channel trials at the end of the previous session (dotted lines). Lines indicate mean across trials and participants and shaded region indicates the standard error of the mean across participants on the five initial trials of each session. Force profiles were aligned to movement start, peak velocity and movement end, prior to averaging. **B**. Retention of the learned force profile (% of learned profile at the end of the previous session). The percent retention was calculated for each participant individually before averaging across participants to calculate the mean. Error bars indicate the standard error of the mean across participants.

### Spontaneous Recovery

After novel dynamics are learned, the introduction of the opposite dynamics (de-adaptation) followed by an error-clamp phase exhibits spontaneous recovery ^27,32^. This rebound to the originally learned motor memory is thought to arise as the de-adaptation removes the fast-learning process, allowing the unveiling of the slow-learning process or processes. Here we investigated whether further exposure to the curl force field results in changes in the extent of spontaneous recovery, by introducing a de-adaptation-error clamp phase in weeks 1, 7 and 23. In the exposure phase prior to the introduction, the kinematic error was close to zero (Fig 7A, blue traces), and the force compensation was between 80% and 100% of full adaptation (Fig 7B, blue traces). The introduction of the opposite curl force field introduced very large negative kinematic errors (Fig 7A, red traces) and resulted in a rapid decrease in force compensation measured on the channel trials to below zero (Fig 7B, red traces). Upon the introduction of the error-clamp phase (consistent channel trials), the predictive force compensation was initially negative, but gradually increased back to compensating for the initially learnt force field direction, plateauing around trials thirty to forty in this error-clamp phase. This spontaneous recovery was larger in week 7 and 23 than in week 1. Finally, the participants were re-exposed to the original curl force field dynamics (re-exposure phase, dark blue traces). While initial kinematic error was large in the positive direction (Fig 7A), this reduced within only a few trials. Similarly, the predictive force compensation recovered very rapidly towards the original levels (Fig 7B). To confirm the presence of spontaneous recovery we used a repeated measures ANOVA to compare the first channel trial after the de-adaptation phase to the mean of the final ten channel trials (main effect phase) for each of the three different weeks (main effect week). Although there was no significant effect of the week (F_2,18_=1.132; p=0.344; 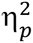=0.112), there was a significant effect of phase (F_1,9_=117.790; p<0.001; 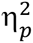=0.929) showing spontaneous recovery, and a significant interaction effect of week and phase (F_2,18_=7.757; p=0.004; 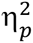=0.463).

**Figure 7.**
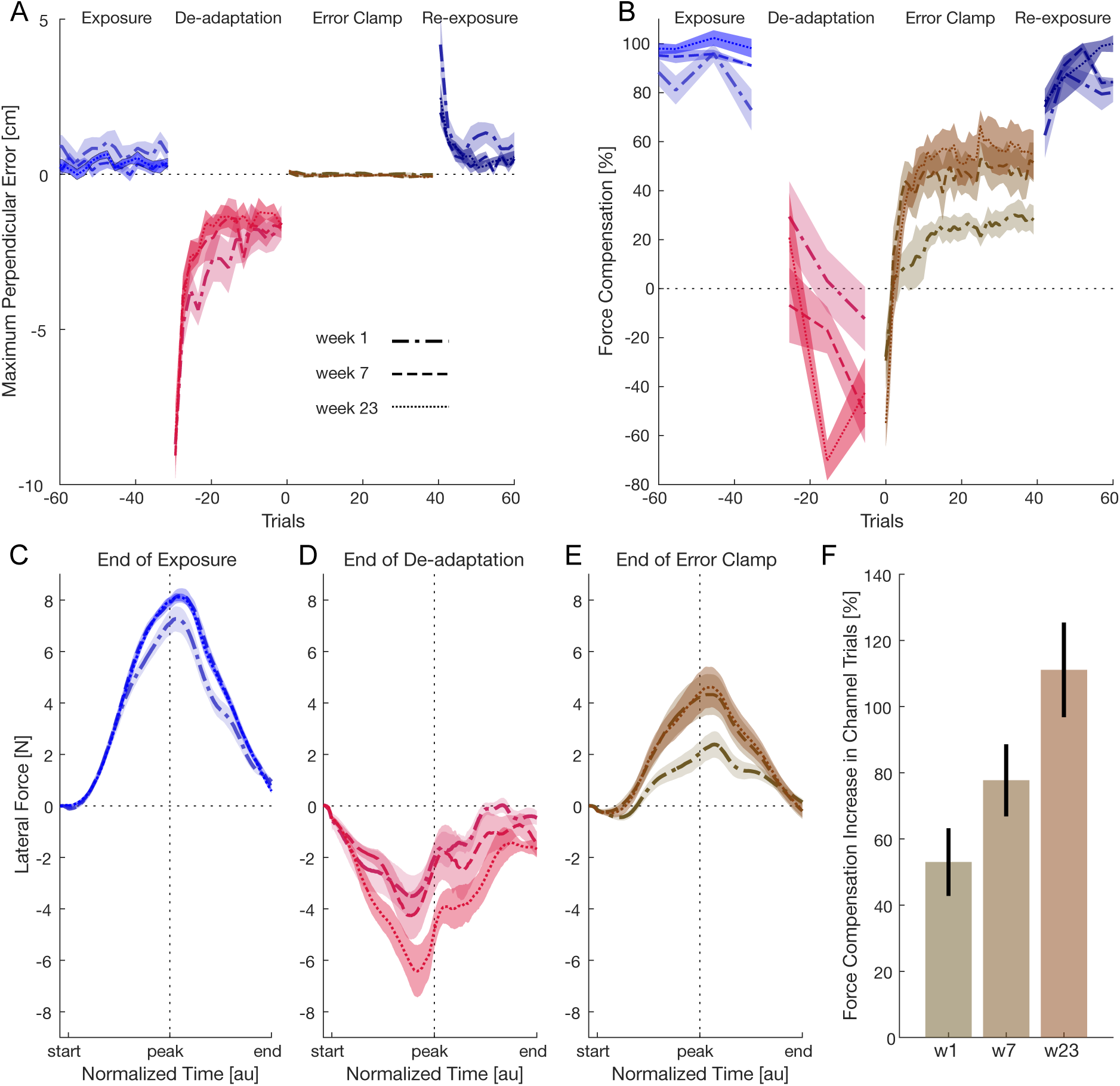
Effect of training on magnitude of spontaneous recovery. **A**. Maximum perpendicular error before, during and after the de-adaptation error clamp phase to measure spontaneous recovery. Positive error reflects perturbations to the left whereas negative errors reflect errors to the right. Colors indicate the phase of the experiment (blue: exposure; red: de-adaptation; brown: error-clamp; dark blue: re-exposure) for weeks 1 (dash-dotted), 7 (dashed) and 23 (dotted). Lines represent the mean across participants and shaded areas represent the standard error of the mean. Trials are aligned to the start of the error-clamp (mechanical channel) phase. **B**. Force compensation before, during and after the de-adaptation error clamp phase. **C.** Lateral force on the final channel trial in the exposure phase before the introduction of the de-adaptation phase. Force profiles were aligned to movement start, peak velocity and movement end, prior to averaging. Shaded region indicates standard error of the mean across participants. **D.** Lateral force on the channel wall at the end of the de-adaptation phase. Value is taken from the very first channel trial in the error-clamp phase. **E.** Lateral force on the final channel trial in the error-clamp phase. **F.** Force compensation increase between the first channel trial and the last channel trial in the error-clamp phase. Error bars indicate standard error of the mean across participants.

To further examine the spontaneous recovery, we plotted the force profiles against the channel walls at the different phases (Fig 7C-E). During the exposure phase, the force profiles showed clear predictive force, with a bell-shaped profile similar to the reaching velocity as expected (Fig 7C). However, by the end of the de-adaptation phase, the force profiles were negative, showing rapid adaptation to the opposite force field, and clear velocity-dependance (Fig 7D). This was particularly enhanced in week 23. However, by the end of the error-clamp phase the rebound towards the original force field was clearly apparent (Fig 7E). Moreover, the force profile aligns well with the velocity profile, with the peak of the forces around the time of peak velocity, and an overall shape that is similar to the original exposure phase force profiles. We find a very strong spontaneous recovery, especially in weeks 7 and 23. Finally, to compare the full extent of the change in force compensation during the error-clamp phase we calculated the difference between the first and last channel trials in this phase (Fig 7F). A repeated measures ANOVA found a significant effect of weeks (F_2,18_=7.757; p=0.004; 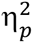=0.463). We find the largest change in week 23 and the smallest change in week 1 (post-hoc comparison found a significant difference between these two weeks (p=0.011). Overall, we find clear evidence for spontaneous recovery, and show that this increases with further training, such that by week 23 the final levels in the channel trials show predictive force compensation close to 60% perfect force adaptation. Interestingly, we also find the strongest de-adaptation in week 23 (Fig 7D) resulting in the largest increase during the error-clamp phase (Fig 7F).

The final level of spontaneous recovery was much larger than seen in previous work, with the levels approaching 60% by week 23. The levels were close to those of the final levels found during the decay and the initial levels for retention measured each day. This suggests a potential connection between these variables, which we examined using linear regression. There were only low correlations between the level of retention and spontaneous recovery (r^2^=0.162; t_78_=3.8865, p=0.0002) or between the final level of decay and the level of retention (r^2^=0.293; t_28_=3.4066, p=0.0020). However, there was a strong correlation between the final levels of spontaneous recovery and final level of decay (r^2^ = 0.657; t_28_=7.3224, p<0.0001). Although the final level of spontaneous recovery varied over the three testing sessions, the final level of decay varied similarly (Fig 8). This suggests a potential connection between both measures and may indicate that both measure a more permanent component of the motor memory.

**Figure 8.**
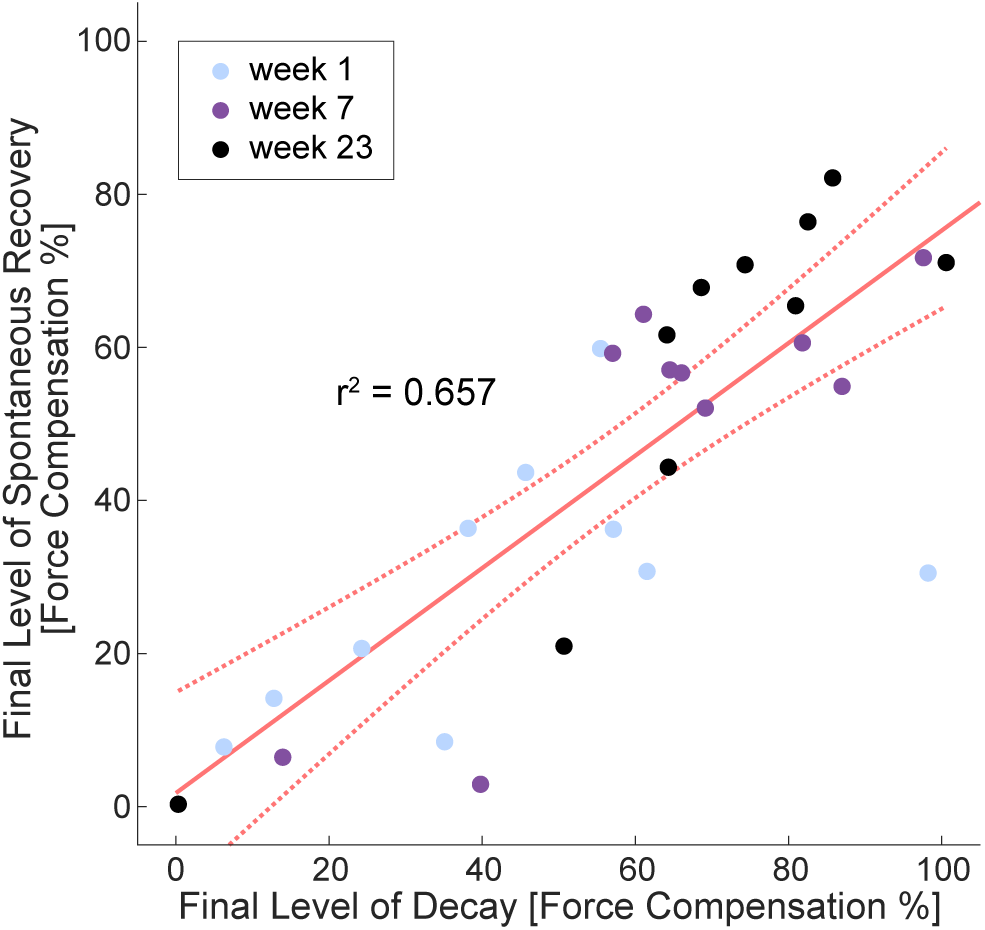
Correlation between spontaneous recovery and final level of decay. The final level of spontaneous recovery for each participant in weeks 1, 7 and 23 is plotted against the final level of decay on the same session. Each point represents the mean force compensation on the last 5 channel trials in the error clamp phase for one participant. Colors indicate the specific session. Line is the best-fit linear relation and dotted lines represent the 95% confidence intervals of the fit.

### Motor Memory Transfer across Limbs

Prior work has examined the transfer of a learned motor memory of a force field to the opposite arm ^47–51^. While the results have suggested very modest transfer of the learning to the opposite arm, around 10%, most of these did not measure the extent of transfer using channel trials. Here we trained participants over thousands of trials with their dominant right hand and examined the transfer of the learning to their left hand in weeks 1, 4, 7 and in the final test on week 23. All left-hand trials to examine the transfer occurred in a mechanical channel allowing us to measure the exact force profile. As movements with the left hand may exhibit forces even in the absence of transfer, we first trained participants to make the reaching movements in the null field on the first session and then measured the baseline forces in channel trials prior to them experiencing the force field.

The force compensation during left-hand movements in the baseline phase were close to zero as expected (Fig 9A) with variability from one participant to another. This was true for both the first baseline session in which participants were always moving with their left-hand (Fig 9A, dark green bar) and when random trials were performed with their left-hand while most trials were right-handed (Fig 9A, light green bar). After the introduction of the force field, the transfer as measured by force compensation (Fig 9A, purple bars) was increased relative to the baseline but only by a small amount. Individual results were variable, and one participant showed a very strong transfer on week 1, which never re-appeared. We can only speculate that this may have resulted from some cognitive strategy, although this is not possible to confirm. On average the force compensation was around 10% across the training sessions. However, a repeated measures ANOVA of the amount of force compensation across the 6 measurement periods found no significant effect of the different measurement periods (F_1.581,14.225_=2.216; p=0.152; 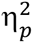=0.198). To further examine the transfer, we plotted the lateral force profile (scaled to the percent level of compensation). Here, the direction of 100% force compensation would indicate a Cartesian transfer of the motor memory while the −100% force compensation would indicate an intrinsic transfer. Importantly, participants never experienced any force field during movements with their left hand. While the forces in the baseline were close to zero, the forces throughout the adaptation period were low but generally positive (Fig 9B). By subtracting the mean baseline force profile, we could look at the exact pattern of forces that were transferred to the left-hand (Fig 9C). These started at zero at the beginning of the movement, peaking just before the time of peak velocity. While there was some variability between sessions, in general the magnitude was around 10% of the full compensation. Finally, to examine individual subjects, we averaged all baseline trials together and all force field trials together to see whether each individual transferred in a similar manner to the left hand (Fig 9D). One possible explanation for the low transfer could have been that some participants transferred in joint coordinates (producing negative force compensation) and others in Cartesian coordinates (producing positive force compensation) resulting in a low average. However, all participants appeared to produce either zero or low but positive force compensation values suggesting what transfer occurred was within a Cartesian framework (Fig 9D). To quantify the transfer, we calculated the mean force compensation change from that in the null field for every participant between the start and end of the movement. The mean of the force compensation across the four tests of transfers (w1, w4, w7, and w23) was positive for all ten participants and significantly different from zero (t_9_=4.338; p=0.002; d=1.372). Overall, we find low levels of transfer of the learned motor memory to the left hand, similar to previous studies. Moreover, the transfer of forces is not well-shaped to the velocity profile of the hand (Fig 9 C,D).

**Figure 9.**
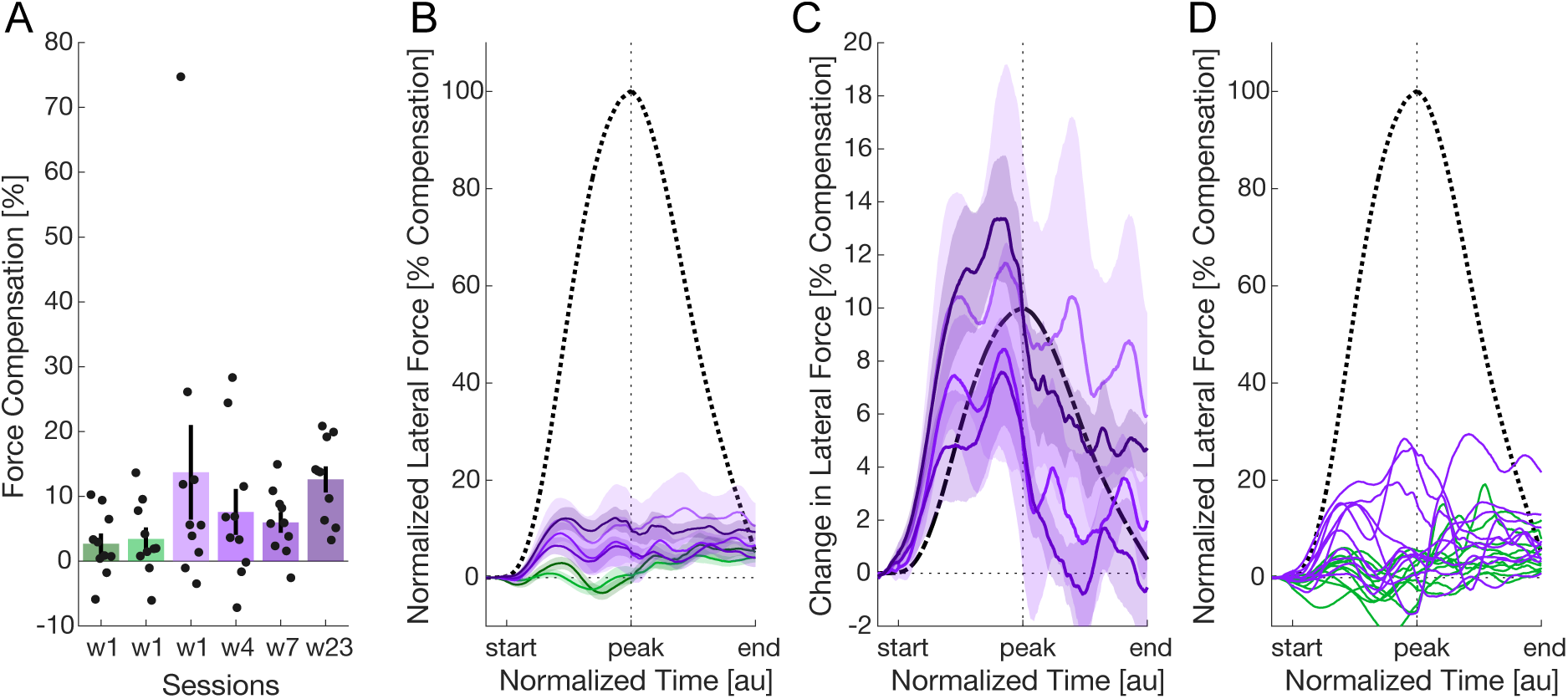
Transfer of the motor memory to the opposite arm. **A**. Force compensation in the left-hand as measured in the baseline condition (green bars) and in weeks 1, 4, 7 and 23 (purple bars). Error bars represent standard error of the mean. Black dots represent individual participants mean values during each session. The left baseline measure (darker green) is performed in all left-hand movement trials. The right baseline measure (lighter green) shows the measure during the occasional left-hand channel trials performed within primarily right-handed movements. **B**. Force profiles of the lateral force against the channel wall for left-hand movements. Solid lines and shaded regions represent the mean and standard error of the mean across all participants in each session. The dotted line represents the force profile that would perfectly compensate for the force field. Force profiles have been aligned to the movement start, peak velocity and movement end prior to averaging. **C**. Change in the force profile after adaptation. The mean force profiles across both baseline sessions were subtracted from the force profiles of each session during the adaptation phases. Dashed line shows the 10% adaptation to the force field if the temporal adaptation was perfectly matched to the velocity of the movement. **D**. The mean baseline force profiles (green lines) and transferred force profiles (purple lines) across all day’s results for each participant. Each line represents one participant. The dotted line represents the force profile that would perfectly compensate for the force field.

## Discussion

This study investigated the long-term changes which occur as a motor memory is formed. Ten participants adapted to a velocity dependent force field applied to a single forward movement for thousands of trials over an eight-week period, followed up 15 weeks later by a retention session. While participants adapted rapidly to the force field in the first session, further weeks of training produced higher compensation, approaching 95% of perfect compensation to the dynamics. While this resulted in straighter and straighter movements, the S-shape of the trajectory was retained throughout the entire experiment. Moreover, retention between weeks increased drastically after week 2. Although de-adaptation appeared to occur faster in the later sessions, the amount of spontaneous recovery was much higher suggesting that the motor memory was more resistant to interference. Despite these strong effects, transfer to the left arm remained very low. Furthermore, we found a higher and narrower angular generalization suggesting changes in the tuning of the neural basis function underlying the motor memory. Finally, the retention session revealed that the learnt motor memory stays well intact despite a 15-week period with no training.

Many previous studies have measured the predictive compensation to novel velocity dependent force fields, finding that force compensation rapidly increases with experience, before gradually plateauing around 80% of perfect adaptation ^10,13,32,40,52–55^. Indeed, it has been suggested that this under compensation is actually an optimal solution to the learning problem ^40^. However, in our study we find that while the initial learning reaches a similar level rapidly on the first day, further experience in the force field drives the predictive force compensation over 95%. This suggests that the previously seen adaptation of only 80% might not reflect the final level of adaptation but only an early stage in dynamic adaptation. With motor memory consolidation the skill acquired becomes more refined ^56–58^. However, as the measure of force compensation only represents the rough level and not the specific spatiotemporal profile of adaptation, we examined the pattern of force produced by the participants against the channel wall and compared it directly to the forces experienced in the force field. In the initial adaptation sessions, the predictive force profiles roughly match the velocity-dependent nature of the force field. However, the pattern of the force profiles tunes further over repeated sessions until it matches the field forces almost perfectly in terms of shape, timing and magnitude. This shows that force field adaptation, just like skill learning ^33–36,58^, continues to improve with further and further practice and highlights the similarity between this novel dynamic task adaptation and these studies of real-world skill learning. Finally, we also found a stabilization of the peak movement velocity through the sessions, where the participants learned to produce peak velocities almost perfectly within the “great” velocity feedback range. The participants learned to perform in accordance with the specific feedback of our experiment on what was considered ‘good’ and ‘great’.

As in all previous studies, the kinematic error reduced with force field exposure. However, the kinematic error continued to reduce even after the first week session, with smaller errors and a reduction in the variability across participants. In addition to the kinematic error reduction, the hand trajectories move in an S-shape towards the end of the adaptation ^28,40,59^, with participants overcompensating for the force fields early in the movement and allowing the force field to bring them back to the target. Izawa and colleagues ^40^ found that even though the paths became straighter with training over three days, the S-shape was retained. They compared these trajectories with an optimal control model, showing that these models predict both the amount (around 80% adaptation) and shape of the trajectories – with overcompensation at the beginning and under-compensation at the end of the trajectory. This S-shape trajectory was also consistently observed throughout the experiments, with the feature still observable at week 8 and during the recall session on week 23. Interestingly, although this S-shape magnitude reduced further with additional training sessions, it was still present even during the final sessions. The presence of the S-shape was originally interpreted as an optimal solution to the problem in which only partial compensation to the novel dynamics occurs ^40^ – thereby reducing the actual compensation and effort needed to still reach the endpoint. However, we find this trajectory present even once the participants have almost perfectly adapted to the force field (week 8) where force compensation values are over 95%. It is a question whether such trajectories are still optimal in this same sense ^40^, or whether after many weeks producing such curved patterns of movement, the final trajectory exhibits this pattern more due to use dependent learning ^60^.

One key finding of the dynamic motor adaptation paradigm is that the memory of the learned task is retained over weeks and months ^22,56,61–63^. However, most of the measures of retention examine how well people move in the force field in subsequent sessions either with reduced hand-paths ^61,62^ or through the correlation of velocity profiles ^22,56,63^. However, neither of these measures can accurately assess the exact level and tuning of the retention, and would be affected by limb stiffness ^11,45^. Moreover, as we can rapidly learn in a few trials or even within one trial ^64,65^, the measures themselves may be influenced by learning occurring within the initial movements. To avoid these issues, we directly assessed the retention across sessions by always applying a mechanical channel in the first five trials in every new session. Although the second session only showed low levels of retention (18.71%), the level of retention was quickly increased to around 60% and was little affected in later weeks. Even in the recall session, 15 weeks after the last training session, participants showed a level of retention around 60%. The development of the retention appears slower than the overall learning of the force field, however the values achieved well matched the final levels of the predictive force compensation at the end of the decay period, suggesting that it might be related to a more permanent component of the memory.

Our motor memories decay when feedback is withheld or the error is clamped to zero ^28,32,41,66–68^. However, it is unclear whether this decay changes with extensive training. It might be expected that, as the memory is stabilized, the extent and rate of decay would reduce, potentially due to the slower timescales of memory (slow or hyper-slow processes) that have higher retention rates ^27,32,44^. Here we probed decay within each training session to investigate potential changes with further adaptation. Interestingly, although the starting level was higher through the sessions, as in previous studies ^59,69,70^, we found little changes in the decay rate or decay extent after the first four weeks of training. Previous studies observed a reduction in decay rates in later training stages ^59,70,71^, which have suggested to indicate the stability of a motor memory ^59^. Their findings could explain why we see subtle reductions in the decay rate between the early training sessions. However, contrary to these studies, we observed a plateau in the decay rate for later weeks, likely due to the length of our study.

Decay has been suggested to be a feature of error-dependent learning which produces trial-to-trial forgetting once feedback is withheld or clamped to zero ^32,41,72^. However, motor adaptation has also been suggested to be a minimization of a cost function balancing error and effort ^7,42,73^. Here, the presence of decay is important for continually searching for an optimal solution that results in lower muscle activation and effort costs. This may partially account for the observed decay in motor memories during error-clamp trials, even when the motor memory appears to be more stable in later training stages. In essence, our findings support the hypothesis of a stabilization of the learnt motor memory over time which seems to plateau after the third or fourth session (corresponding to around 3000 training trials). While we can only speculate about whether there is a connection or not, the level of plateau in the decay matched very well the level of spontaneous recovery between sessions. This level might reflect the portion of the motor memory which is resistant to trial-to-trial or temporal decay ^74^.

While spontaneous recovery in cognitive tasks has a long standing history ^75,76^, the phenomenon has also been observed in motor learning studies ^18,27,32,77–80^, where it has been used to support the existence of multiple adaptation processes. The theory is that there is both fast and slow states of learning, and the spontaneous recovery reflects the level of the slow state ^32,80^. Here we consistently observed spontaneous recovery (weeks 1, 7, and 23) with significant differences between weeks. The level of spontaneous recovery increased from around 20% in week 1 to approximately 60% in week 23. This finding aligns with previous research, showing increased spontaneous recovery after longer exposure in both word association ^81^ and motor learning paradigms ^82,83^. They suggested that increased recovery does not stem from a higher learning rate of the slow process but rather from a higher retention of the slow process. Our experiment extends these findings by demonstrating increased spontaneous recovery after prolonged exposure to original learning by implementing this paradigm in different weeks in our experiment. We propose that the substantial increase observed in spontaneous recovery over training weeks supports our hypothesis of a more stable motor memory following extended exposure.

One important question in motor adaptation is the transfer of the learned skill to different effectors. There is evidence that motor memories of force field adaptation can be transferred to the other limb, although this transfer appears modest. This transfer occurs both from the dominant hand to the non-dominant hand and vice versa ^47,49^. However, the amount that is transferred decreases as the time after the initial learning occurs ^49^. While there has been some suggestion based on the larger intermanual transfer for abruptly introduced dynamics compared to gradually introduced dynamics that the transfer may be primarily explicit in nature ^84^, it appears that the transfer is stronger and lasts longer with longer training schedules ^48^. In general the amount of intermanual transfer is small, with estimates using channel trials of around 9-12% ^48^. In our study, we also found similarly low levels of transfer. However, we found little to no differences across weeks 1, 4, 7 or 23. This may be because we already have extensive training on week 1 compared to previous studies ^48,84^. Importantly, the shape of the transferred force profile does not match the velocity profile, suggesting the transfer was limited and not well tuned to the force field dynamics. As the left hand never directly experienced the force field, the transfer likely occurred either from the visual experience of the right hand being perturbed during the training sessions ^85^ or directly from the experience of the forces (e.g. due to shared neural representations ^16,26,86,87^. While we also found this transfer was in Cartesian coordinates, this pattern can be explained for both the visual ^85^ and dynamic errors ^88^. Overall, we found clear, but limited transfer to the left hand in a Cartesian reference frame which was unaffected by the long-term training.

When learning a new motor skill, we build motor memories of the external world which also generalize to similar reaching angle directions ^12,19,28,89–96^. The motor memory decays away from the learned reaching direction resulting in local angular generalization which is well modelled by a Gaussian function ^31,90,93,94,97^. We also found a Gaussian-like angular generalization which slightly narrowed with increased exposure suggesting a slight fine tuning of the spatial properties of the motor memory over the training weeks. Researchers have hypothesized that neural tuning functions underlie this angular generalization ^28,89,98^, and that the fast and slow processes of motor adaptation might have distinct tuning properties ^99,100^. They found, similar to our study, that the generalization slightly decreased with learning ^100^. However, one might also expect a decrease in the angular generalization independent of such changes in the fast and slow processes. It has been suggested that when we suddenly experience a change in the tasks, leading to the formation of a new motor memory, that the motor memory initially shows a wide generalization which is gradually tuned and narrowed towards the specific new task ^2^. Our results show the clear narrowing of the generalization function, although we are unable to determine whether this might be related to changes in the which basis functions represent the motor memory ^100^ or occurring due to narrower tuning of the neural basis functions themselves ^2^.

Our work clearly shows that there continues to be changes in the development of a motor memory even after a few thousand trials. Importantly this is true even for our task – a simple forward reaching in a single direction in a curl force field. This has important implications for a large body of research in the field of motor control, which often studies adaptation using a few hundred trials at most. Our work shows large changes in the level of retention and decay which required around 2000 trials before there was any stabilization of the motor memory. Despite this, there continued to be more subtle changes, even between week 7 and the testing on week 23, in the measures of spontaneous recovery. While subtle, it appeared that in week 23 participants adapted to the opposite force field much quicker and still showed strong spontaneous recovery. These long-term differences in the motor memory development have very important implications for rehabilitation.

## Methods

### Experimental Participants

Ten young adults participated in this series of experiments (3 male and 7 female: aged 26.9 ± 2.0, mean ± SD). All participants were force field naïve. All participants participated in this experiment once every week for 8 weeks (5-9 days apart) and a final 9^th^ session that took place 15 weeks after the 8^th^ session (week 23). Each session lasted between one (e.g. session 3) and two hours (session 1), corresponding to around 10 hours of training over 9 sessions for each participant. All participants were right-handed according to the Edinburgh handedness inventory ^101^ with no reported neurological disorders. The study was approved by the Ethics Committee of the Medical Faculty of the Technical University of Munich. All participants provided a written informed consent before participating in the study.

### Apparatus

Participants, seated in a chair, grasped the handle of the vBOT robotic manipulandum ^102^ with both of their forearms supported against gravity with air sleds (Fig. 10A). The robotic manipulandum was used to both generate the environmental dynamics (null field, force field or channel), and measure the participants’ behavior. Position and force data were sampled at 1kHz. Endpoint forces at the handle were measured using an ATI Nano 25 6-axis force-torque transducer (ATI Industrial Automation, NC, USA). The position of the vBOT handle was calculated from joint-position sensors (58SA; IED) on the motor axes. Visual feedback was provided using a computer monitor mounted above the vBOT and projected veridically to the participant via a mirror. This virtual reality system covers the manipulandum, arm and hand of the participant. Participants performed forward reaching movements in the horizontal plane at approximately 15 cm below the participants’ shoulder level. These were mainly performed with the right hand.

**Figure 10.**
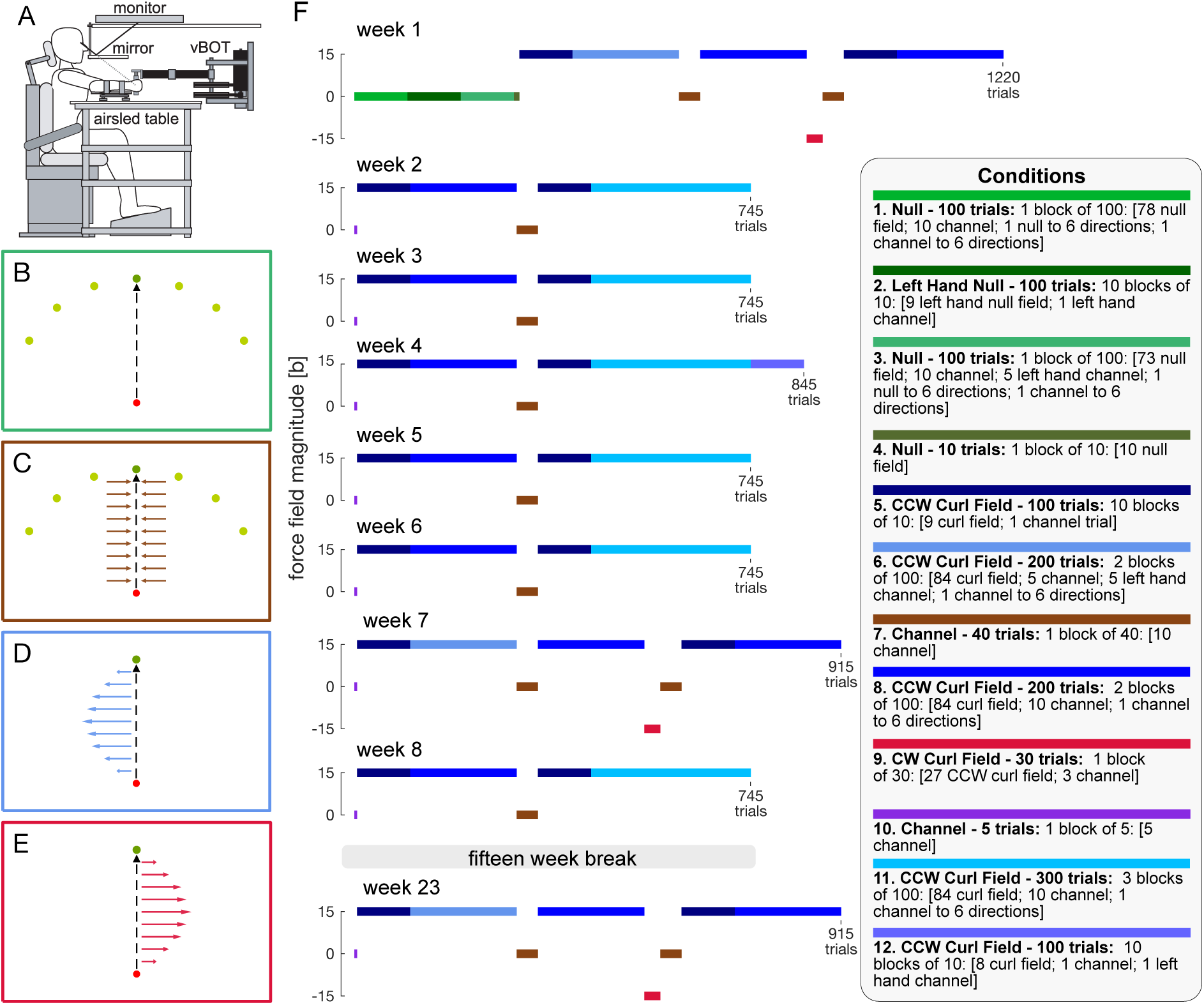
Experimental Design. **A.** Participants were seated and grasped the handle of a robotic manipulandum. Visual feedback about the task and the location of their hand was provided through a monitor reflected by a mirror such that it appeared in the plane of movement. The arm was supported by an airsled on a table. **B.** In the baseline session (week 1) movements were performed from the start position (red circle) to one of 7 targets (green circles) in the null field. **C.** Throughout the entire experiment, random trials could be performed in a mechanical channel (shown here for the straight-ahead target). These occurred to all seven possible target directions. **D.** Adaptation was examined to a CCW curl force field throughout all weeks. The force field was only ever presented for the target directly ahead of the start position and only for movements with the right hand. **E.** To assess spontaneous recovery, on weeks 1, 7 and 23, the CW curl force field was applied for 30 trials to de-adapt the motor memory. This was only presented for the target directly ahead of the start position on right hand movement. **F.** The detailed protocol of the experimental sessions. Colors indicate specific blocks as detailed on in the legend of conditions. Within a block, all trials were pseudorandomized. In the legend, the color of each condition is shown directly above the description of the specific trials.

### Experimental Setup

Participants were seated with their shoulders restrained against the back of a chair by a shoulder harness (Fig 10A). Movements were made from a 1.4 cm diameter start circle centered approximately 25.0 cm in front of the participant to a 1.6 cm diameter target circle centered 25 cm in front of the start circle. The participant’s arms were hidden from view by the virtual reality visual system, on which the start and target circles as well as a 1.0 cm diameter cursor used to track instantaneous hand position were projected. Participants were instructed to perform successful movements to complete the experiment. A successful movement required the hand cursor to enter the target (without overshooting) with a peak speed of 60 ± 8 cm/s of movement initiation. Overshoot was defined as movements that exceeded the target in the direction of movement. When participants performed successful movements, they were provided with feedback as to how close they were to the ideal peak speed of 60 cm/s (‘great’ if the speed was within ± 4 cm/s of the desired peak speed, otherwise they received ‘good’ feedback) and the counter increased. Similarly, when they performed unsuccessful movements, they were provided with feedback as to why the movement was not considered successful (“too fast”, “too slow” or “overshot target”). Trials were self-paced; participants initiated a trial by moving the hand cursor into the start circle and holding it within the target for 1000 ms. A beep then indicated that the participants could begin the movement to the target. The duration of the movement was determined from the time that the participants exited the starting position until the time that participants entered the target. Once in the target, they were required to stay within the target for 1000 ms before the trial ended. At the end of each trial, the vBOT passively moved the participant’s hand to the starting location, using a minimum jerk trajectory. Rest breaks were provided after each 200 trials. Participants were also instructed that they could take a break anytime between trials by simply removing the cursor from the start target. Almost all movements required the participant to use their right hand, however on occasional trials the left-hand movement was also assessed.

### Force fields

In the adaptation movement, participants performed reaching movements either in a null field condition (Fig 10B), a velocity-dependent curl force field or a mechanical channel (Fig 10C). The curl force fields were implemented as:

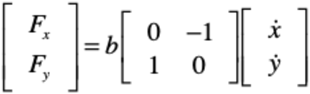

where the field constant b was set to a value of ±15 Nm^-1^s, and the sign determines the direction (CW or CCW) of the force-field. All participants experienced the same force field direction (CCW) as their main learning protocol (Fig 10D), but the opposite force field (CW) was provided in three sessions as part of a spontaneous recovery measurement paradigm (Fig 10E). Mechanical channel trials ^41,103^ were implemented using a spring constant of 6,000 Nm^-1^ and a damping constant of 10 Nm^-1^s perpendicular to the direction of motion throughout the movement between the central location and the final target.

### Experimental design

The goal of this experiment was to examine motor adaptation to a simple force field for a single direction of movement over a long period of training. Therefore, our study required each participant to do one training session each week for 8 consecutive weeks (+/- 2 days) followed by a recall session 15 weeks after the last training day (week 23). In addition to examining the changes in the kinematics and predictive force compensation of the learned movement, our experimental design also assessed the sharpness of the tuning of the generalization, interlimb transfer, decay, retention of the motor memory across training sessions and the degree of spontaneous recovery using a de-adaptation-error clamp protocol.

#### Generalization Tuning

All participants were presented with the force field only when reaching directly forward to a target (0° target). However, we know that this motor memory generalizes to nearby reaching angles ^28,89,90,104,105^. To assess the tuning width of the generalization of the learned motor memory to nearby directions, participants made occasional reaching movements to six other targets angled −60°, − 40°, −20°, 20°, 40° and 60° away from the 0° target. The distance from the starting circle was 25cm for all movements. In the pre-exposure phase on week 1, these movements could be in the null field (Fig 10B) or a channel trial (Fig 10C). However, once the force field was presented to the participants, all angled trials were in the mechanical channel. These angled trials were presented during every experimental session.

#### Interlimb Transfer

All participants were only presented with the force field when reaching with their right hand. It has been suggested that motor memories can be transferred to the other limb after adaptation ^47–49,106,107^ although there is some suggestion that this is driven more by explicit strategies ^84^. To test possible interlimb transfer and how this might evolve over multiple training sessions, we examined the transfer to left hand reaching movements during weeks 1, 4, 7 and 23. In the first week, prior to force field exposure, participants made reaching movements with their left hand in the null force field (90 trials) and in a mechanical channel (15 trials). After the introduction of the force field, left hand movements were always in a mechanical channel. Prior to each left hand movement, the screen instructed participants to switch the robot to their left hand. Once they released the handle and grasped the handle with their left hand, the robot passively moved their left hand to the starting position. A similar process occurred when they needed to switch back to the right hand. All movements with right or left hands were performed using the same robotic handle.

#### Motor Memory Decay

Multiple studies have shown that adaptation is a process of both adaptation to error and retention of the existing motor memory ^7,32,41,73^. That is, that at least two parameters (learning rate and retention rate) are needed to explain adaptation ^41^. More recently many studies have suggested that adaptation may consist of multiple learning and retention rates, one for each process ^27,32,43,77^. To determine whether the decay of a motor memory when no error is presented changes with longer learning, we introduced 40 trials of mechanical channels (error clamp trials) in a row after a long period of experiencing the force field. This was repeated on every session of the experiment (9 times in total) to examine whether the decay is reduced (higher retention term) with longer training over multiple days of exposure.

#### Retention of Motor Memory between Sessions

It has long been well known that the learned motor memory on one day is retained over hours, days, or even months of rest, and can be quickly selected and used when faced with the same force fields upon returning to the laboratory ^22,56,61,108^. However, all studies generally apply the force field immediately on the first few trials on the subsequent days, making it difficult to determine exactly how much of the learned motor memory is retained and actively used in the absence of error information. To assess the amount of retained motor memory, we introduced 5 channel trials in a row at the beginning of each session after the force field was introduced (weeks 2-23). This could be compared across sessions and against the final predictive force compensation on the previous session.

#### Spontaneous Recovery

The presence of spontaneous recovery in an error-clamp phase after the presentation of de-adaptation trials has supported the idea of multiple timescales of motor memory formation ^27,32,52^. Here we applied 30 trials of a CW force field followed by 40 trials within a mechanical channel on week 1, 7 and 23.

### Protocol

In order to look at the learning effect over multiple sessions of a force field adaptation task, we set up the following protocol (Fig 10F). All trials were done with their dominant (right) hand, except where specified.

#### Week 1

Participants started with a baseline session (310 trials total) in the null field with their right and left hands (Fig 1F), including channel trials and movements to different target locations (+/- 20°, 40° and 60° angles). After this baseline session, a CCW velocity dependent curl force field was introduced for 300 trials including 10% channel trials, followed by consecutive 40 channel trials to measure decay. After another 200 trials in the force field, the opposite curl force field (CW) was applied for 30 trials followed by 40 channel trials in a row to assess spontaneous recovery. This was followed by a further 300 trials in the CCW curl force field. During the CCW force field training, left hand assessment (only in channel) and channel trials in all target directions were interspersed randomly. The experimental session was 1220 trials in total (see Fig 10F for specific details).

#### Weeks 2, 3, 5, 6 and 8

To start each experimental session, participants first moved in 5 channel trials in a row, followed by 300 reaching movements in the CCW curl force field (10% of channel trials randomly applied in a block design). This was followed by 40 consecutive channel trials to measure motor memory decay, and then 400 trials in the CCW curl field with random channel trials (all movement directions). Each experimental session was comprised of 745 trials total (Fig 10F).

#### Week 4

After starting with 5 consecutive channel trials, participants made 300 reaching movements in the CCW curl force field (10% channel trials). This was followed by 40 consecutive channel trials to measure decay. This was followed by 400 trials in the CCW field with channel trials in all directions. Finally, 100 trials were performed in the CCW with random tests of transfer to the left hand (performed in channel trials). The experimental session in week 4 was a total of 845 trials (Fig 10F).

#### Weeks 7 and 23

After starting the experiment with 5 consecutive channel trials, participants made 300 reaching movements in the curl force field (10% of channel trials). This was followed by 40 consecutive channel trials to measure decay. After another 200 trials in the force field, the opposite curl force field (CW) was applied for 30 trials followed by 40 channel trials in a row to assess spontaneous recovery. This was followed by a further 300 trials in the CCW curl force field, containing left hand channel trials and random channel trials in all target directions. The experimental sessions in weeks 7 and 23 were 915 trials in total (Fig 10F).

### Analysis

The data was analyzed using Matlab R2024b. Force and kinematic data were low-pass filtered at 40Hz with a fifth-order, zero phase-lag Butterworth filter. Individual trials were aligned on movement onset. For each trial, we calculated measures of kinematic error or force compensation between 200 ms prior to leaving the start position until 200 ms after entering the target position.

For each non-channel trial, the absolute maximum perpendicular error (MPE) was calculated and used as a measure of the kinematic error. The MPE is the absolute maximum perpendicular distance between the hand trajectory and the straight line between the start and end targets. For each channel trial, the force compensation was calculated as a measure of the predictive compensation to the CCW force field. The force compensation is calculated by the regression between the force produced by participants into the wall of the simulated channel (lateral measured force) and the force needed to compensate perfectly for the force field. Here the perfect compensatory endpoint force is determined on each trial using the forward velocity on each trial, and the magnitude of the force field. Force compensation was always calculated with respect to full compensation in the CCW curl force field (indicated as 100% force compensation) such that perfect adaptation in the CW force field would produce −100% adaptation.

#### Force Traces

To examine the pattern of force adaptation to the dynamics, the force traces were calculated across specific channel trials. As the force applied by the force field is velocity dependent, and each trial could have a different duration, in order to average across multiple trials, the force traces were aligned to the start of the movement, the time of peak velocity and the end of the movement. Here the start and end of the movement were defined as the time at which the velocity first and last exceeded 5% of the peak velocity on each trial, respectively. Both the first half (start to peak velocity) and the second half (peak velocity to movement end) were resampled to 300 points. Individual trials from the same participant or across different individuals could then be averaged together without affecting the shape of the force profiles. For an accurate comparison of the exact forces applied to the robotic handle in the force field, the same procedure was used in the curl force field trials once the field had been learned to plot the actual forces experienced.

#### Motor Memory Generalization

To examine the force compensation generalization to nearby target angles, the mean force compensation for each target direction [−60°, - 40°, −20°, 0°, 20°, 40°, 60°] was calculated for each participant on each week’s training. This data was then fit by a Gaussian function centered at 0° to determine the height and width of the Gaussian. For each training week, we fit the all-possible combinations of eight of the ten participants data to obtain a set of 45 parameters. For each parameter fit, we ran the optimization (fminsearch) 2000 times from random parameter starting positions and selected the parameter for the best-fit. Results were plotted using the boxchart function in Matlab allowing us to compare how the parameters changed over the weeks of training using the 95% confidence intervals of the parameter estimates.

#### Motor Memory Decay

To fit the time constant of the decay in channel trials, an exponential function was fit the force compensation over the 40 consecutive channel trials to obtain the time constant and magnitude of the decay. The initial force field compensation was determined using the 5 last channel trials in the CCW prior to the introduction of the error-clamp phase. For each training week, we fit the all-possible combinations of eight of the ten participants data to obtain a set of 45 parameters. For each parameter fit, we ran the optimization (fminsearchbnd) 2000 times from random parameter starting positions and selected the parameter for the best-fit. Results were plotted using the boxchart function in Matlab allowing us to compare how the parameters changed over the weeks of training using the 95% confidence intervals of the parameter estimates.

#### Statistics

Statistics were performed in JASP 0.19.2 ^109^. Statistical significance was considered at the p<0.05 level. For repeated measures ANOVAs, a Mauchly test was applied to check for sphericity. In the case that the Mauchly test was significant, the degrees of freedom were adjusted using a Greenhouse-Geisser correction. We report 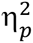 as the effect size and used Bonferroni post-hoc tests to determine any differences between levels. For t-tests, we tested normal distribution via the Shapiro-Wilk test and presented the effect size as Cohen’s d.

